# PromoterAtlas: decoding regulatory sequences across Gammaproteobacteria using a transformer model

**DOI:** 10.1101/2025.07.11.664312

**Authors:** Lucas Coppens, Rodrigo Ledesma-Amaro

## Abstract

Recent advances in deep learning, particularly transformer architectures, have improved computational approaches for biological sequence analysis. Despite these advances, computational models for bacterial promoter prediction have remained limited by small datasets, species-specific training, and binary classification approaches rather than comprehensive annotation frameworks. We present PromoterAtlas, a 1.8M parameter transformer model trained on 9M regulatory sequences from 3,371 gammaproteobacterial species. The model demonstrates recognition of various regulatory elements across different species, including ribosomal binding sites, various types of bacterial promoters, transcription factor binding sites, and terminators. Using this model, we developed a whole-genome promoter annotation tool for Gammaproteobacteria, with various levels of validation that support the predictions of promoters associated with different sigma (σ) factors. Furthermore, we show that the model embeddings encode cross-species evolutionary relationships, clustering promoters by σ factor identity rather than species-specific sequence features. Finally, we show that model embeddings encode regulatory sequence information that enables effective prediction of transcription and translation levels. PromoterAtlas can contribute to our understanding of and ability to engineer bacterial regulatory sequences with potential applications in bacterial biology, synthetic biology, and comparative genomics.

## Introduction

Sequence-based transformer models have recently revolutionised bioinformatics. In the genomic domain, massive models such as Enformer^1^, Nucleotide Transformer^2^, Evo 2^3^, and DNABert-2^4^ have demonstrated unprecedented capabilities in capturing complex sequence patterns and functional relationships across DNA. In the protein domain, the model ESM-2 and the associated ESMFold in particular have shown that purely sequence-based models can be used to predict structural properties with remarkable accuracy, enabling a breakthrough in protein folding^5^. These models leverage self-attention mechanisms to identify long-range patterns between nucleotides or amino acids. By learning contextual representations directly from genomic or protein sequences, transformer architectures have managed to overcome the limitations of previous architectures and approaches that relied heavily on hand-crafted features or position-specific scoring matrices.

Bacterial promoter prediction has been addressed through various computational approaches, including position weight matrices (PWMs), hidden Markov models, support vector machines, convolutional neural networks, recurrent neural networks, and most recently also transformer architectures^6–15^. Deep learning approaches have struggled to improve prediction accuracy over traditional methods like PWMs and HMMs, as they remain constrained by several key limitations^16^. The scarcity of experimentally validated promoters creates a bottleneck in training data, with most models relying on small sets of positive labels that inadequately represent the diversity of bacterial regulatory elements. Furthermore, bacterial promoter models are often trained on data limited to single species, most often *Escherichia coli*, limiting both dataset size and model generalisability. Perhaps most critically, existing approaches typically focus on classifying isolated sequences rather than providing a comprehensive annotation framework. This methodological gap has left automated whole-genome promoter annotation an unresolved challenge, hampering our ability to rapidly annotate newly sequenced bacterial genomes and characterise regulatory landscapes.

We devised an approach incorporating multiple solutions to address the above constraints in bacterial promoter prediction. First, by leveraging unsupervised masked-token learning, we circumvent the limitation of scarce labelled data, allowing the model to discover intrinsic patterns within sequences without requiring explicit promoter annotations. We strategically create a focused dataset consisting of regions spanning 200 nucleotides upstream of coding sequences, thereby creating an enrichment for relevant regulatory signals while reducing noise from irrelevant genomic regions. Finally, we extracted data from a wide range of sequenced genomes of species across the clade of Gammaproteobacteria, capitalising on presumed evolutionary conservation of core promoter architecture within this phylogenetic group. The Gammaproteobacteria is one of the most extensively studied bacterial clades, spanning approximately 250 genera and containing diverse medically and biotechnologically relevant species such as *E. coli, Pseudomonas aeruginosa, Pseudomonas putida, Salmonella enterica, Vibrio cholerae, Vibrio natriegens, Yersinia pestis*, and *Klebsiella pneumoniae*. This taxonomic breadth introduces sufficient sequence diversity to capture variations in promoter structures while maintaining conservation of core regulatory mechanisms.

In this paper we present PromoterAtlas, a 1.8 million parameter model trained on approximately 9 million regulatory sequences from 3,371 diverse gammaproteobacterial species. The model employs a “DNATransformer” architecture that combines convolutional filters for capturing local sequence patterns with rotary attention layers for detecting long-range dependencies, pre-trained using masked-token prediction. The base model enables visual identification of functional regulatory elements including promoters, ribosomal binding sites (RBS), transcription factor binding sites, and terminators. We extend this capability by developing a segmentation head that transforms embeddings from the base model into annotations for genome-wide identification of promoters across five σ factors in Gammaproteobacteria, with validation through ChIP-seq data comparison, analysis of predicted promoter distributions, and gene ontology enrichment studies. Additionally, we demonstrate that the model captures evolutionary relationships between different promoter types across phylogenetically distant species, as revealed through analyses of its embedding space. Finally, we show that PromoterAtlas embeddings can be leveraged for sequence-based prediction of gene expression at both transcription and translation levels.

## Results

The PromoterAtlas-base model architecture consists of a linear encoder to project one-hot encoded DNA sequences from dimension 4 to hidden dimension 128, followed by 8 DNATransformer blocks, and finally a linear decoder to project back to dimension 4 (Figure 1A, B). Our DNATransformer block architecture was designed to incorporate both local pattern recognition through a double convolutional filter as well as longer-range relationships through a rotary attention block, each combined with residual connections. Attention with rotary position embeddings was chosen instead of traditional position encodings as it is more suitable for relative position modelling compared to traditional attention with positional encodings, and it has been shown to work well for DNA sequences^17^. The third element of the DNATransformer block is a feed forward block consisting of a linear projection to dimension 256 followed by a linear projection back to dimension 128, again with residual connection. The PromoterAtlas-base model was trained on approximately 9 million 200 nucleotide sequences found upstream of annotated coding sequences in the genomes of 3371 gammaproteobacterial species (Figure 1C). The machine learning task was masked token prediction on zero-masked positions. In every 200-nucleotide sequence, 20 random positions (10%) were masked. The core concept is that to accurately predict the missing positions, the transformer model must learn to recognise and use contextual sequence information. This encourages the model to focus on regularly reoccurring patterns in the datasets, such as promoter motifs, as these are more predictable than less biologically relevant positions. Training converged after 264 epochs and showed good generalisation between training and validation dataset, which represented a 9:1 split of the data (Supplementary Figure S1, Materials and methods).

**Figure 1.**
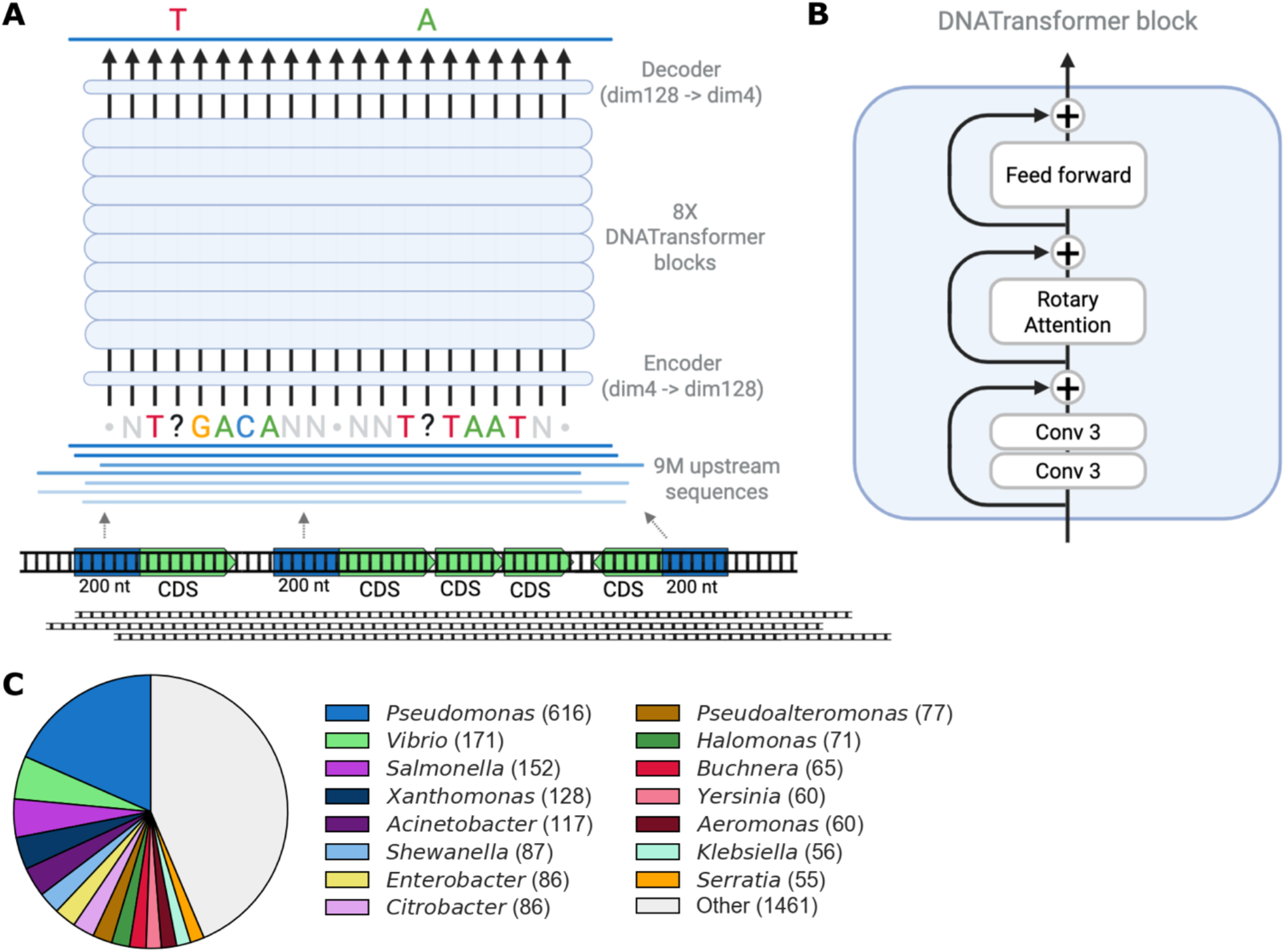
Pre-training a base model on regulatory sequences across the clade of the Gammaproteobacteria. (A) Overview of the pre-training workflow. 200 nucleotide sequences upstream of all coding sequences (CDS) are extracted from the genomes of the Gammaproteobacteria in the training set. During training, positions are randomly masked, and the transformer model is optimised to predict the missing tokens from the sequence’s context, encouraging it to learn relevant sequence features such as promoter motifs (e.g. TTGACA-TATAAT). (B) Overview of the DNATransformer block architecture used in PromoterAtlas. (C) Overview of Gammaproteobacterial genera included in the data set used to pre-train PromoterAtlas.

### Recognition of regulatory features

Plotting the per-base pre-softmax logits after the linear projection in the decoder provided a good way of visualising the sequence patterns recognised by the pre-trained model in individual sequences (Figure 2). This visualisation method revealed a wide variety of recognisable sequence features including promoters corresponding to at least 5 σ factors as well as RBS. Palindromic terminators were also often found to be recognised by the model despite only sequences upstream of coding sequences having been included in the training dataset. This reflects the underlying genomic architecture where sequences 200 nucleotides upstream of coding sequences in the dataset often included terminators from the previous genes due to the relatively short distances between bacterial genes. Similarly, the logit plots showed that PromoterAtlas recognised stop codons when the preceding gene overlapped with the analysed 200 nucleotide upstream sequences. We also found that the model showed recognition of known motifs of various transcription factor binding sites, including motifs for CRP, ntrC, luxR, and FNR (Supplementary Figure S2).

**Figure 2.**
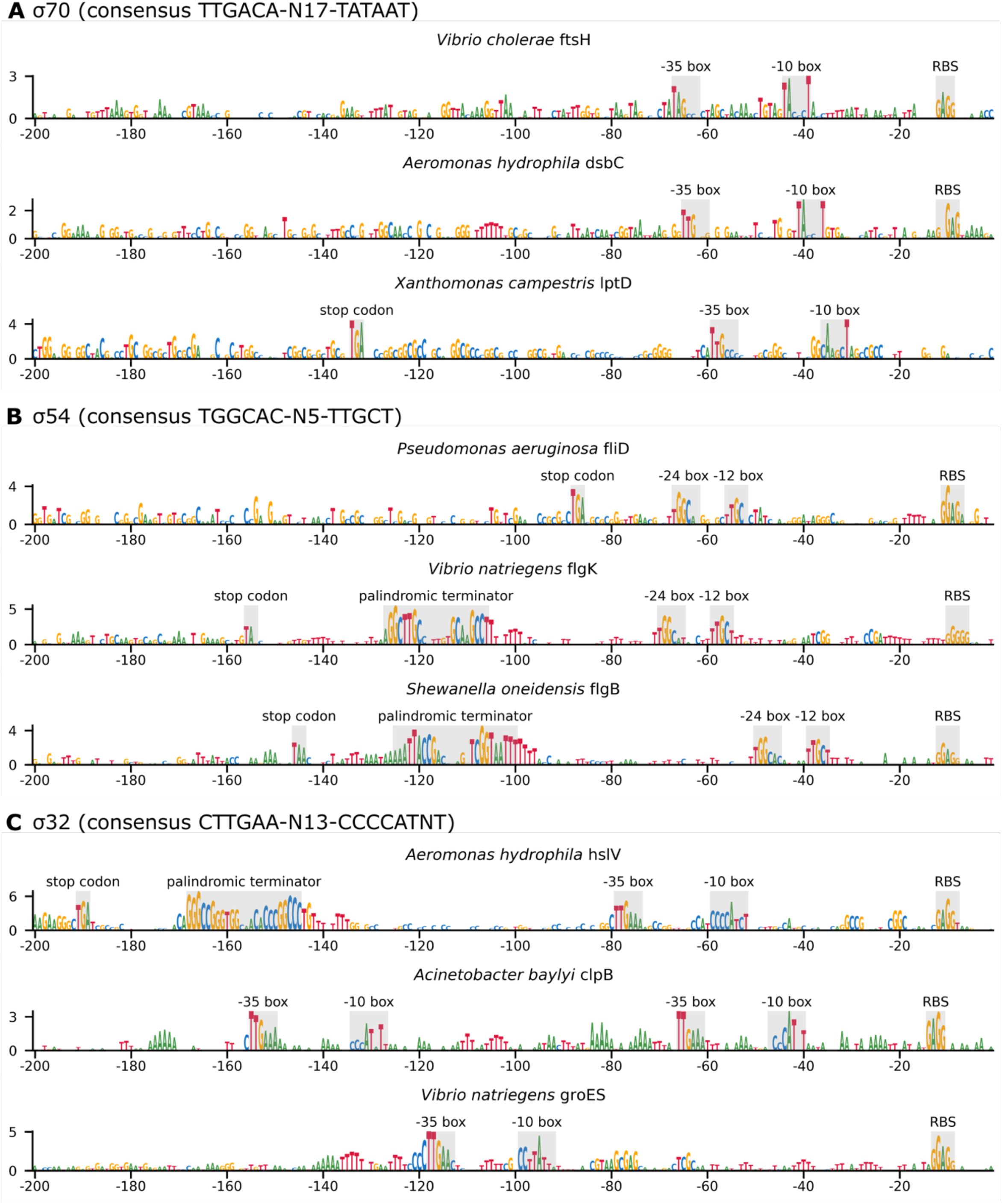
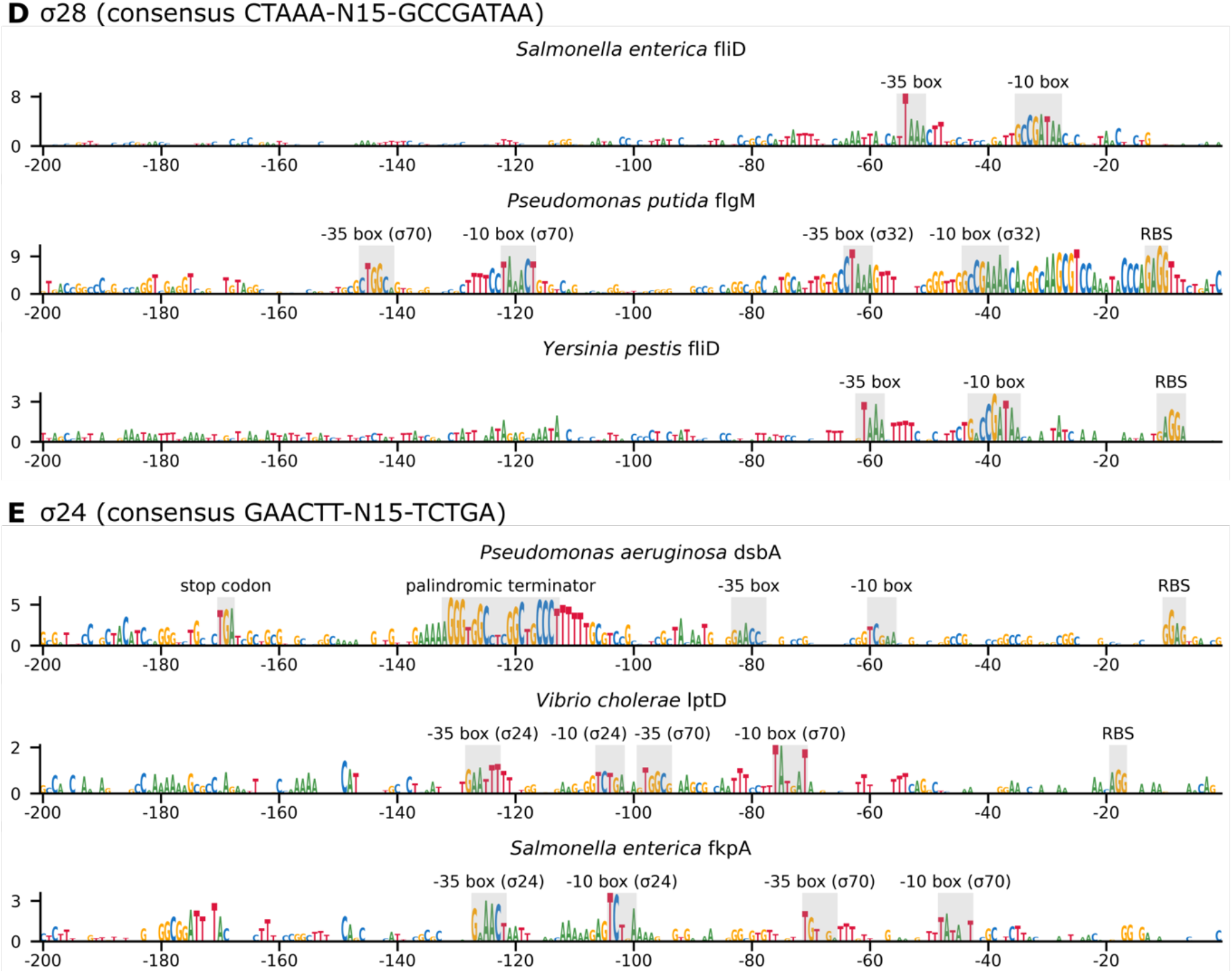
Logit plots of regulatory sequences upstream of coding sequences show recognition of a wide variety of motifs in various Gammaproteobacteria including RBS, motifs of promoters belonging to at least five σ factors, and terminators. The manually added annotations show what features are being recognised by PromoterAtlas in the areas with high per-nucleotide logits (A) logit plots showing recognition of σ70 promoter motifs (B) logit plots showing recognition of σ54 promoter motifs (C) logit plots showing recognition of σ32 promoter motifs (D) logit plots showing recognition of σ28 promoter motifs (E) logit plots showing recognition of σ24 promoter motifs.

As the PromoterAtlas logit plots highlighted a large diversity of features in each of the tested species, these per-base logit plots may serve as a novel and useful tool for regulatory sequence analysis in Gammaproteobacteria. In some cases, particularly for upstream regions of genes known to have high expression levels in bacteria, large logit values along long stretches of the analysed sequences impeded the clear distinction between promoter motifs and surrounding nucleotides (Supplementary Figure S3). In those cases, we found that plotting the transformer’s attention maps was helpful in revealing the location of the promoter motifs. These attention maps also revealed that the model focuses especially on the final position in the −10 box of the promoters.

### Whole genome promoter motif annotation

Since the PromoterAtlas-base logit plots of many tested sequences highlighted sequence features that we could easily recognise as known promoter patterns (Figure 2), we decided to make a tool that could automatically recognise these patterns and annotate them as promoter motifs. To do this we trained a segmentation head that can transform the PromoterAtlas-base embeddings to a per-position discrete signal that can be programmatically converted to an annotation (Figure 3A). The training data for this segmentation head was a set of 200 nucleotide sequences with obviously recognisable promoter motifs that we manually annotated based on logit plots, consisting of 147 σ70/σ38 promoters, 50 σ54 promoters, 76 σ32 promoters, 27 σ28 promoters, and 18 σ24 promoters. The σ70 and σ38 promoter motifs were grouped together for the segmentation task as they are too similar to visually distinguish between, given their often overlapping affinity for both σ factor polymerase complexes^18^.

**Figure 3.**
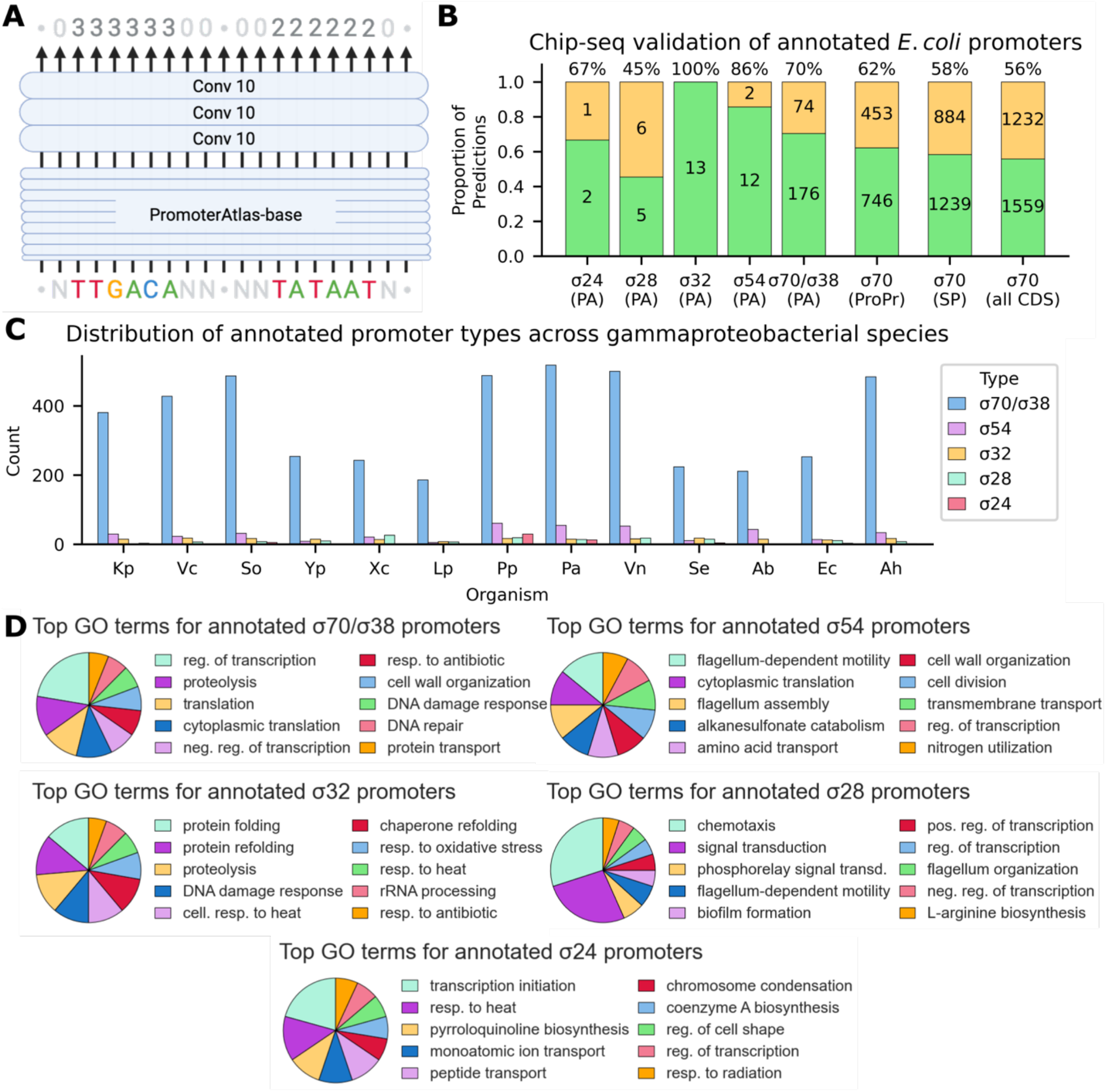
A segmentation head leveraging the PromoterAtlas-base embedding enables automated genome-wide promoter annotation for 5 σ factors. (A) A segmentation head consisting of three convolutional filters of width 10 allows to transform the PromoterAtlas-base final layer embeddings into per-position discrete outputs that can be used for automated sequence annotation. (B) Overlap between the *E. coli* ChIP-seq data from Cho et al. (2014)^19^ and PromoterAtlas (PA) annotations, ProPr and SAPPHIRE (SP) σ70 promoter predictions, and a naive baseline scenario in which σ70 promoters are automatically predicted 30 nucleotides in front of every coding sequence in the *E. coli* genome. (C) The number of automatically annotated promoter motifs for each σ factor in selected Gammaproteobacteria: Kp (*K. pneumonia*), Vc (*V. cholerae*), So (*Shewanella oneidensis)*, Yp (*Y. pestis*), Xc (*Xanthomonas campestris*), Lp (*L. pneumophila*), Pp (*P. putida*), Pa (*P. aeruginosa*), Vn (*V. natriegens*), Se (*S. enterica*), Ab (*Acinetobacter baylyli*), Ec (*E. coli*), Ah (*Aeromonas hydrophila*). (D) The top 10 gene ontology terms associated with the genes following predicted annotations of the different σ factors in (C).

Critically, no promoters were annotated for training of the segmentation head in regulatory sequences from *E. coli*, as this was the only organism for which we found genome-wide Chip-seq data for various σ factor binding sites^19^, presenting an opportunity to validate the segmentation tool on out-of-training data. The results of this validation experiment with the trained PromoterAtlas-annotation model are shown in Figure 3B. The majority of predicted promoters fell within the genomic ranges identified by Cho *et al*. (2014)^19^’s ChIP-seq experiment, reaching up to a 100% accuracy in the case of the 13 predicted σ32 promoters. Crucially, for each of the predicted but unvalidated σ24, σ28, and σ54 promoters, as well as each of the analysed σ70, we found that the logit plots exhibited clear promoter motifs matching the binding sites of these σ factors (Supplementary Figure S4). We therefore deemed these predictions plausible, particularly considering that ChIP-seq experiments are inherently condition-dependent and unlikely to have captured the complete set of functional promoters across the diverse environmental and physiological states that *E. coli* can undergo.

We also evaluated PromoterAtlas-annotation against existing tools by comparing its σ70 promoter predictions with those from SAPPHIRE^9^ and ProPr^20^. While more widely cited tools like BPROM^6^ and CNNPromoter^11^ exist, their web interfaces allow maximally one DNA sequence to be analysed at a time and therefore do not support high-throughput analysis of regulatory sequences extracted from genbank files, highlighting the need for end-to-end annotation pipelines like ours. When applying the same ChIP-seq validation criteria, both ProPr and SAPPHIRE predicted substantially more putative σ70 promoters but with lower precision (Figure 3B). Notably, SAPPHIRE and ProPr achieved validation rates of 58% and 62% respectively, compared to 56% for a naive baseline scenario in which σ70 promoters are systematically placed 30 nucleotides upstream of every coding sequence, indicating modest improvement over this uninformed positional heuristic. In contrast, PromoterAtlas-annotation achieved a 70% validation rate, indicating higher specificity through fewer false positives, though, as noted earlier, the validation rates plausibly underestimate actual performance based on the presence of clear promoter motifs in the ‘false positives’ and the condition-dependent limitations of ChIP-seq data. While this conservative approach, achieved by training exclusively on sequences with obviously recognisable promoter motifs, inevitably results in lower sensitivity due to promoters with less obvious motifs being missed, it aims to ensure that the annotations provided are of high confidence and correspond to clearly recognisable regulatory elements suitable for reliable downstream analyses. Furthermore, our approach extends beyond existing tools by providing annotation capabilities for four additional σ factors, representing a functional advantage over current methods.

We subsequently used the segmentation model to rapidly predict and annotate hundreds of promoter motifs in a variety of medically and biotechnologically relevant gammaproteobacterial species (Figure 3C). PromoterAtlas-annotation annotates varying amounts of promoters depending on the species, with the smallest amount of promoters predicted in *Legionella pneumophila* (206), and the largest amount of promoters detected in *P. aeruginosa* (615). This may suggest variability in the level of conservation of promoter motifs between species, though the high representation of *Pseudomonas* species in the pre-training dataset (Figure 1C) might also partially account for the model’s enhanced pattern recognition within the *Pseudomonas* genus. The relative abundances of promoters predicted by PromoterAtlas-annotation across different σ factors are plausible from a biological perspective. For instance, the proportion of predicted σ70/σ38:σ54 promoters is roughly 12:1, which more closely reflects the natural distribution by which these promoters are known to occur in bacteria than the 3:1 ratio present in the manually annotated promoter data used to train the segmentation model. Similar divergences from training data proportions are evident when examining the other σ factors as well, with the model consistently predicting distributions that align with biological expectations. This suggests that PromoterAtlas-annotation is recognising genuine promoter patterns rather than merely reflecting the data bias of its training set.

Finally, we examined whether the genes downstream of the annotated promoter motifs correspond to biological functions typically associated with the respective σ factors. Figure 3D presents the top ten gene ontology (GO) terms enriched in genes positioned downstream of predicted promoters across the species shown in Figure 3C. The σ70/σ38-driven genes primarily relate to fundamental cellular processes including regulation of transcription and translation, along with their regulation, reflecting σ70’s role as the housekeeping σ factor^21^. The σ54-regulated genes show enrichment in functions relating to flagellar motility and nitrogen utilisation, representing known σ54-related functions^22^. Genes downstream of σ32 promoters are strongly associated with protein folding, protein refolding, chaperone functions, and heat response pathways, matching its canonical role in heat shock response^23^. The σ28-regulated genes also demonstrate clear enrichment in known associated roles such as chemotaxis, signal transduction, and flagellum-related functions^24^. Finally, genes controlled by σ24 promoters show associations with heat response, ion transport, and peptide transport, consistent with its function in regulating extracytoplasmic stress responses^25^. This consistency between predicted promoter annotations and expected biological functions provides further validation of PromoterAtlas-annotation accuracy in identifying genuine σ factor binding sites.

### PromoterAtlas encodes cross-species evolutionary relationships

Given PromoterAtlas’s ability to identify promoter motifs belonging to different σ factors in diverse species spanning the Gammaproteobacteria, we investigated whether the model has a notion of the shared evolutionary origin of the different σ factors between various species. To explore this question, we extracted sequence embeddings from various layers of the PromoterAtlas-base model for promoters from multiple organisms and σ factor types. For each sequence, we applied global max pooling across the sequence length to create a fixed-length vector representation that captures the most salient features detected by the model. These sequence-level embeddings were then visualised using Uniform Manifold Approximation and Projection (UMAP)^26^ to reveal clustering patterns across species and σ factor types.

Analysis of embeddings across different network layers (after DNATransformer blocks 1, 3, 5, and 7) revealed a progressive shift in the model’s representation space. In the shallow layers (particularly after block 1), the primary organising principle appeared to be genomic GC content, with high-GC organisms *P. aerugiosa* and *X. campestris* forming a distinct cluster from a lower-GC cluster with *E. coli* and *V. natriegens*, regardless of promoter type (Figure 4). However, as information is passed through to deeper layers, σ factor identity emerged as the dominant organising principle. By block 7, promoters from the same σ factor class clustered together across species boundaries, with distinct groupings observed for σ54, σ32, and σ28 promoters. This hierarchical organisation of the PromoterAtlas-base embedding space suggests that the model has captured the shared evolutionary conservation of σ factor binding motifs across Gammaproteobacterial species. The transition from GC content to σ factor-based organisation may reflect the evolutionary history of these regulatory elements, where core promoter recognition mechanisms likely predate species-specific GC content biases.

**Figure 4.**
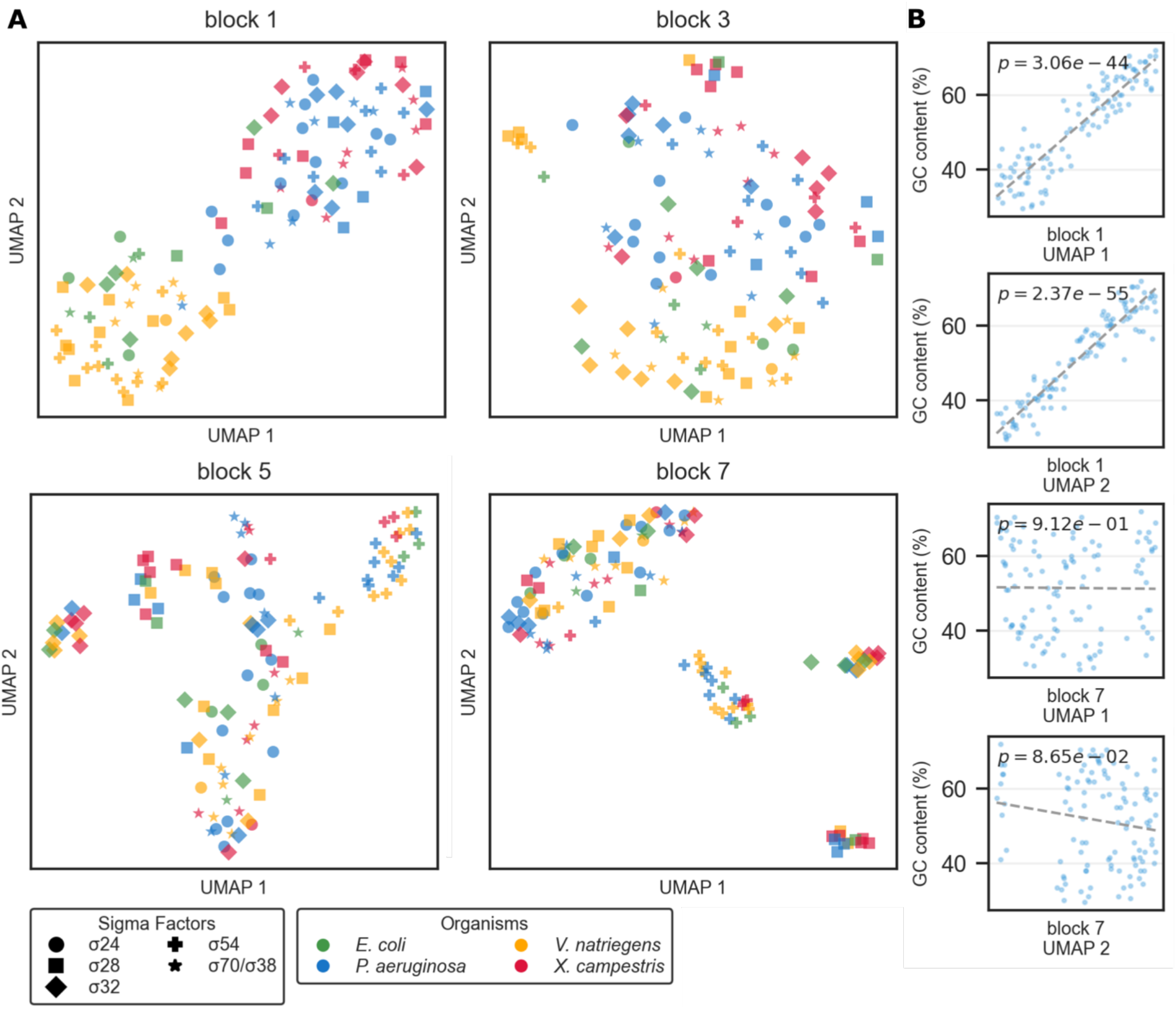
Representations of the embedding space of various σ type promoters across *E. coli, V. natriegens, P. aeruginosa*, and *X. campestris*. (A) UMAP dimensionality reduction plots of the pooled embeddings of various σ type promoters extracted from the sequence representations following DNATransformer block 1, 3, 5, and 7. (B) Scatter plots of sequence GC content against UMAP dimensions 1 and 2 from block 1 and block 7. The UMAP dimensions show strong correlation with GC content in the representations following block 1 while the correlation between UMAP dimensions and GC content is not significant in the representations following block 7.

Interestingly, in the visualisation of the block 7 embedding space, some instances of σ54, σ32, and σ28 promoters were clustered with the main heterogeneous central grouping rather than their σ-specific smaller clusters. To investigate this phenomenon, we examined the positions in the sequences that most strongly contributed to the pooled sequence embeddings. For promoters within the distinct σ factor clusters, we identified a consistent pattern where specific positions in the embedding space were consistently activated regardless of where the characteristic motifs appeared in the input sequence (shown for σ28 in Supplementary Figure S5). This suggests that the model employs a positional encoding mechanism to group functionally related promoters. However, the biological basis for why some promoters of a given σ type are represented through this mechanism while others are not remains unclear. This divergence might reflect subtle functional distinctions within σ factor categories that are not captured by conventional motif-based classification. While this observation provides insight into the computational representation strategy employed by the model, a complete biological interpretation of these different encoding patterns requires further investigation.

### Predicting transcription and translation levels

To evaluate PromoterAtlas’s ability to capture functional regulatory information beyond motif identification, we assessed its performance in predicting gene expression at both transcription and translation levels.

For the transcription prediction task, we used datasets from multiple sources: Lafleur et al. (2022)^27^ (5,392 sequences), Hossain et al. (2020)^28^ (4,350 sequences), Urtecho et al. (2019)^29^ (10,898 sequences), and Yu et al. (2021)^30^ (1,494 sequences). All sequences were standardised to a maximum length of 86 nucleotides with shorter sequences padded with adenine nucleotides. Two prediction models were implemented to compare the effectiveness of learned embeddings versus direct sequence encoding. The primary model leveraged the pre-trained PromoterAtlas-base backbone with frozen weights to extract feature embeddings (Figure 5A). These embeddings were then fed into a prediction head consisting of two 1D convolutional layers (kernel size 8, stride 1, with ‘same’ padding) followed by max pooling (kernel size 7, stride 1, padding 3), maintaining the base model’s hidden dimension of 128. The pooled features were flattened and passed through two fully connected layers (dimensions 128×86→64→1) with ReLU activations between layers to predict normalised transcription levels. For comparison, we implemented a simplified model that applied the same convolutional architecture directly to one-hot encoded DNA sequences (4 channels), bypassing the transformer embeddings entirely. This model used identical convolutional parameters, pooling operations, and fully connected layers (dimensions 128×86→64→1) as the primary model, but operated on raw sequence one-hot encodings rather than learned embeddings.

**Figure 5.**
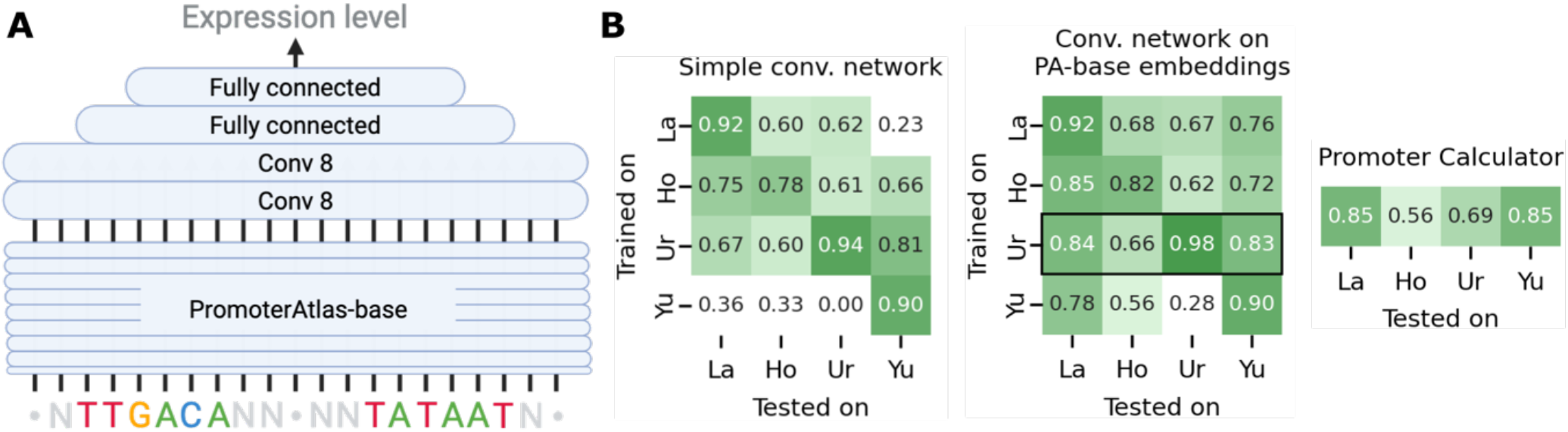
A prediction head leveraging the PromoterAtlas-base embedding enables sequence-based prediction of expression levels (A) A prediction head consisting of two convolutional filters of width 8 followed by pooling and fully connected layers allows to transform the PromoterAtlas-base final layer embeddings into real values that can be used to capture expression levels (B) Heatmaps showing the Pearson correlations between real and predicted transcription levels using various methods trained and tested on four datasets: Lafleur et al (2022)^27^, Hossain et al. (2020)^28^, Urtecho et al. (2019)^29^, Yu et al. (2021)^30^. The predictor that combines PromoterAtlas-base embeddings with the convolutional prediction head consistently outperforms and generalises better across the datasets than the model consisting of just the convolutional network without relying on the PromoterAtlas-base embeddings. The PromoterAtlas-based model trained on the Urtecho et al. (2019)^29^ dataset is highlighted as the best transcription prediction model, with Pearson correlations either on par with or better than the Promoter Calculator.

Both transcription prediction models were trained four times, each time using 80% of one of the four datasets, and using 10% validation data for early stopping. The remaining 10% were used for cross-data testing of each of the resulting models. The results (Figure 5B) demonstrate that the PromoterAtlas-based model consistently outperformed the simple convolutional network across all datasets. This indicates that the PromoterAtlas-base model captures biologically relevant regulatory information enabling more accurate prediction of gene expression. Notably, the best performing model, the PromoterAtlas-based model trained on the Urtecho et al. (2019)^29^ dataset, reached Pearson correlation values on the four datasets which were either on par with or higher than the correlation values we found using Promoter Calculator^27^. This suggests that this model in particular reaches state-of-the-art performance on the task of sequence-based prediction of transcription levels.

We applied an identical methodology for the prediction of protein expression levels accounting for both transcription and translation levels. For this task we used data from Kosuri et al. (2013)^31^, which included combined promoter and RBS sequences with corresponding protein expression measurements. We employed the same architectural design as our transcription prediction model, using the pre-trained PromoterAtlas-base backbone with frozen weights. Using the same train:validate:test split of 8:1:1 for training and early stopping, the trained protein expression model reached a Pearson correlation of 0.79 on the test set (Supplementary Figure S6). This correlation approaches the advanced ANOVA model correlation of 0.82 reported by Kosuri et al. (2013)^31^ but achieves this performance without requiring interaction modelling between RNA and protein levels. Instead, the PromoterAtlas-based model effectively captures the regulatory relationships governing both transcription and translation processes using sequence information alone.

## Discussion & perspectives

In this work, we presented PromoterAtlas, a transformer-based model trained on 9 million regulatory sequences from 3,371 gammaproteobacterial species. We demonstrated that PromoterAtlas successfully recognises diverse regulatory elements including promoters for five different σ factors, RBS, and terminators across the gammaproteobacterial clade. By leveraging the base model embeddings, we developed a genome-wide promoter annotation tool capable of identifying σ factor promoters. Furthermore, we showed that PromoterAtlas embeddings enable state-of-the-art performance in predicting transcription and translation levels from sequence alone, and that the model’s embedding space encodes meaningful relationships between regulatory elements of different bacteria.

A key aspect underpinning this work is the approach to data collection. By systematically extracting upstream sequences of coding regions across a diverse range of species rather than relying on experimentally validated promoters, the model could learn from a much wider array of regulatory contexts than previously possible. This approach might allow the model to capture subtle sequence features that influence DNA physical properties such as melting temperature or topological characteristics. These features are typically overlooked by traditional position weight matrices or other methods exclusively focussed on motifs. Given the vast number of sequence examples it has encountered, the transformer architecture might develop more sophisticated representations of molecular features related to promoter function, potentially identifying permissible and prohibited base combinations that extend beyond our current scientific understanding. Transformer models focussed on specific features and trained across many species within taxonomic clades may find applications in the analysis of other biological functions as well.

Our work presented the first end-to-end GenBank annotation tool specifically designed for bacterial promoters. We took a conservative approach and focussed only on the most clearly recognisable regulatory patterns due to the absence of comprehensive experimental validation data across diverse bacterial species. This method was validated through multiple strategies: ChIP-seq data comparison in *E. coli*, analysis of the relative abundances of predicted σ-type promoters, and gene ontology enrichment studies. While promising, further refinement with additional experimental data stemming from a diverse range of species would likely enhance the model’s performance, as well as open up ways to expand the predictions beyond visually obviously recognisable promoter motifs.

Current expression prediction tools for bacterial systems are primarily developed for and validated in *E. coli*, creating a need for more broadly applicable approaches. While the expression prediction models presented in this work were also trained and validated using *E. coli* datasets due to data availability constraints, PromoterAtlas’s underlying base model was trained across the entire gammaproteobacterial clade. This broader phylogenetic training foundation suggests that our approach may be more readily extensible to expression prediction in non-model bacteria compared to existing tools. This hypothesis is supported by our segmentation model results and UMAP analysis, which demonstrate that the later-layer embeddings encode regulatory information that transcends species boundaries. Nevertheless, comprehensive cross-species validation of expression prediction capabilities remains challenging due to the scarcity of standardised expression datasets for non-model organisms.

Looking forward, PromoterAtlas opens several avenues for future research. The model’s ability to identify diverse σ factor binding sites creates opportunities for strength prediction of promoters across different σ factors, potentially extending to transcription factor-dependent expression prediction. Such tools could enhance the understanding of bacterial gene regulation beyond the standard housekeeping promoters. In synthetic biology, accurate prediction and design of promoter strength across different regulatory contexts could facilitate more precise engineering of bacterial gene expression systems. For basic research, the model offers tools for identifying adaptation strategies through regulatory analysis and suggesting potential biological functions for uncharacterised proteins based on their upstream regulatory elements. The embedding space of PromoterAtlas may also enable novel approaches to cluster analysis for identifying co-regulated genes across diverse bacterial species.

While our results demonstrate the potential of transformer models for bacterial regulatory sequence analysis, the current work represents an early step. Future work should focus on incorporating additional experimental data, extending the model’s capabilities to other clades, and developing more sophisticated tools for regulatory sequence engineering based on the model’s learned representations.

## Methods

### Dataset generation

The dataset used for base model training in this study was created by using the NCBI API to query the NCBI genomes database^32^ for genomes of Gammaproteobacteria with annotated coding sequences. Only bacteria with unique species names were retained to prevent identical or highly similar DNA sequences occurring in the dataset. Sequences spanning 200 nucleotides upstream of any annotated coding sequence were extracted and added to the dataset, with the exception of coding sequences occurring within less than 50 nucleotides from the previous coding sequence to avoid including coding sequences belonging to the same operon and therefore potentially lacking promoter or other regulatory features.

### PromoterAtlas-base architecture

Sequences are one-hot encoded before passing to the PromoterAtlas-base network. The network itself starts with an encoder consisting of a linear projection from the one-hot-encoded dimension of four (A, C, G, T) to a hidden dimension of 128. The resulting representation is subsequently transformed by eight DNATransformer blocks, before being projected back to the dimension four by a linear decoder to enable comparison to the missing nucleotide tokens for loss calculation (Figure 1A). A DNATransformer block consists of three elements, which maintain the sequence length of 200 and each have residual connections. The first element is a double convolution with window size 3. The second element is a rotary attention layer, which is a variation of classical attention where positional information is encoded by rotating the query and key token vectors of the attention operator, in a frequency-dependent manner where different rotation frequencies are applied to different vector dimensions^33^. This makes it encode relative positions between features in the sequence, which is well suited to the nature of DNA regulatory sequences where the distance between motifs is highly relevant to their function. The third element of the DNATransformer block is a feed forward layer, consisting of a linear projection from dimension 128 to dimension 256 and subsequently a linear projection back to dimension 128.

### PromoterAtlas-base training

PromoterAtlas-base training was performed by zero-masking 20 positions randomly selected in every sequence, and making the model predict the missing nucleotides. This encourages the model to focus on patterns in the data that can help to determine what nucleotides were masked. The dataset was split into a training set and validation set in a 9:1 ratio. Categorical cross-entropy loss was applied to the masked positions only as the training criterion, using the AdamW optimiser. A batch size of 1024 and initial learning rate of 10^−3^ were used. The learning rate was scheduled to be reduced by half with a validation loss patience of 5 epochs. Early stopping was scheduled with a patience of 15 on the validation set. PromoterAtlas-base was trained on a single NVIDIA a100, and took 264 epochs (∼5 days) to converge. A training loss plot can be found in Supplementary Figure S1.

### PromoterAtlas-base logit plots

The PromoterAtlas-base logit plots are created by extracting the pre-softmax output logits of an input sequence. For each position in a sequence, the value corresponding to the sequence’s nucleotide in that position is used to plot the height of that nucleotide’s letter in the plot. For the attention map plots, the attention matrices for a sequence are extracted from each of the model’s rotary attention modules and subsequently averaged per position. The per-position per-nucleotide averages are subsequently plotted in analogy to the logit plots.

### PromoterAtlas-annotation

The data used to train PromoterAtlas-annotation consisted of 318 sequences spanning 200 nucleotides upstream of coding sequences, in which we manually annotated motifs clearly identifiable in the sequences’ logit plots and attention maps. This resulted in a dataset consisting of regulatory sequences containing 147 σ70/σ38 promoters, 50 σ54 promoters, 76 σ32 promoters, 27 σ28 promoters, and 18 σ24 promoters.

The segmentation model architecture consisted of a pre-trained PromoterAtlas-base backbone with frozen weights, followed by a custom segmentation head. The feature embeddings from the final layer of the backbone were fed into the segmentation head, which consisted of three sequential 1D convolutional layers (kernel size 10, stride 1, with ‘same’ padding). The final convolutional layer projected the features to a 12-dimensional output space, corresponding to the number of possible sequence annotations at each position (e.g ‘no feature’, ‘σ70/σ38 −10 promoter motif’, ‘σ70/σ38 −35 promoter motif’, etc.). Training was performed using cross-entropy loss and the Adam optimizer with a learning rate of 10^−5^ and a batch size of 5. Model performance was monitored on a validation set (20% of the data), with early stopping implemented using a patience of 10 epochs on validation loss.

For end-to-end whole-genome annotation of genbank files, all 200 nucleotide sequences upstream of coding sequences were automatically extracted. Subsequently, promoter features were predicted using the PromoterAtlas-annotation model. The model identified regulatory elements by applying a threshold-based segmentation approach, requiring a minimum of 4 consecutive nucleotides with the same prediction to constitute a valid segment. Co-occurrence rules were enforced to maintain biological consistency, ensuring paired promoter elements (e.g., −10/-35 motifs) appeared together for each σ factor (σ70/σ38, σ54, σ32, σ28, and σ24). Predicted regulatory features were then mapped back to their genomic coordinates, with strand-specific adjustments, and integrated into the original GenBank files as annotated regulatory elements with appropriate feature qualifiers. This automated pipeline facilitated large-scale annotation of bacterial genomes, enabling systematic identification of promoter features across multiple species.

### Gene ontology analysis

To functionally characterise genes regulated by different σ factors, we performed Gene Ontology (GO) term enrichment analysis on genes downstream of predicted promoter motifs. For each annotated promoter identified by PromoterAtlas-annotation, we extracted the locus tag or gene name of the downstream coding sequence from the GenBank files. We then queried the UniProt REST API (https://rest.uniprot.org) to retrieve the associated GO terms for each gene. For each σ factor category, we compiled the frequency of associated GO terms and identified the top ten most enriched terms.

### Embedding space analysis

We selected a random subset of regulatory sequences classified as promoters belonging to the different σ factors by PromoterAtlas-annotation from *E. coli, V. natriegens, P. aeruginosa*, and *X. campestris*. For each of the sequences, we extracted the embedding from multiple layers of the network (blocks 1, 3, 5, and 7), applying global max pooling across the sequence dimension to create fixed-length vector representations (length equal to the hidden dimension 128 of PromoterAtlas-base). The resulting sequence embeddings were then visualised using Uniform Manifold Approximation and Projection (UMAP)^26^ to reduce dimensionality while preserving local and global structure. UMAP parameters were set to n_neighbors=15, min_dist=0.1, and n_components=2. Before UMAP transformation, embeddings were standardised to ensure equal weighting of each dimension. For Supplementary Figure S5, we identified the positions that yielded the maximum value for each dimension in the embedding space and tallied their frequency. These position counts were then visualised alongside logit plots to reveal potential relationships between sequence features and embedding patterns, using colour intensity to represent the number of dimensions along which each position contributed the maximum value for the global max pooling.

### Prediction of transcription levels

We obtained the datasets for the prediction task of transcription levels of promoter sequences from various sources. From Lafleur *et al*. (2022)^27^ we obtained 5,392 promoter sequences, from Hossain *et al*. (2020)^28^ we obtained 4,350 promoter sequences, from Urtecho *et al*. (2019)^29^ we obtained 10,898 promoter sequences, and from Yu *et al*. (2021)^30^ we obtained 1494 sequences. All sequences were standardised to a maximum length of 86 nucleotides with shorter sequences padded with adenine nucleotides.

Two prediction models were implemented to compare the effectiveness of learned embeddings versus direct sequence encoding (Figure 5). The primary model leveraged the pre-trained PromoterAtlas-base backbone with frozen weights to extract feature embeddings. These embeddings were then fed into a custom prediction head consisting of two 1D convolutional layers (kernel size 8, stride 1, with ‘same’ padding) followed by max pooling (kernel size 7, stride 1, padding 3), maintaining the base model’s hidden dimension of 128. The pooled features were flattened and passed through two fully connected layers (dimensions 128×86→64→1) with ReLU activations between layers to predict normalised transcription levels.

For baseline comparison, we implemented a simplified model that applied the same convolutional architecture directly to one-hot encoded DNA sequences (4 channels), bypassing the transformer embeddings entirely. This model used identical convolutional parameters, pooling operations, and fully connected layers (dimensions 128×86→64→1) as the primary model, but operated on raw sequence one-hot encodings rather than learned embeddings.

Training for both models was performed using the AdamW optimizer with a learning rate of 10^−5^ and batch size of 32. Mean squared error was used as the loss function, with model performance monitored on a validation set (10% of data). Early stopping was implemented with a patience of 10 epochs on validation loss, and the remaining 10% of data was reserved for testing. For cross-evaluation, separate models were trained on each dataset and evaluated against all other datasets to assess generalization capabilities. Performance was measured using Pearson correlation between predicted and observed transcription levels, with both model predictions and PromoterCalculator^27^ predictions compared on the same test sets.

For Promoter Calculator predictions^27^, we retained the minimum predicted dG_total across the forward sequence as the prediction score.

### Prediction of protein expression levels

For the protein expression level prediction task accounting for both transcription and translation, we used data from Kosuri *et al*. (2013)^31^, which included combined promoter and RBS sequences with corresponding protein expression measurements. Following our approach for transcription prediction, we standardized all sequences to a maximum length of 86 nucleotides, with shorter sequences padded with adenine nucleotides. For expression level prediction, we employed the same architectural design as our transcription prediction model, based on the pre-trained PromoterAtlas-base backbone with frozen weights. As with the transcription model, feature embeddings from the backbone were processed through a custom prediction head consisting of two 1D convolutional layers (kernel size 8, stride 1, with ‘same’ padding), max pooling (kernel size 7, stride 1, padding 3), and two fully connected layers (dimensions 128×86→64→1) with ReLU activations.

Given the wide dynamic range of protein expression values in the dataset, we applied a logarithmic transformation (log(count+1)) to the target values before normalising. The training procedure followed the same approach as our transcription prediction models, using the AdamW optimiser, with learning rate of 10^-5 and early stopping patience of 10 epochs. We maintained the same train/validation/test split ratio (80%/10%/10%) and evaluated model performance using Pearson correlation between predicted and observed protein levels after reversing the normalisation and logarithmic transformations.

## Data availability

The data used in this study to train the base model as well as the manually annotated promoter motif data for the segmentation model are available at https://huggingface.co/datasets/LCoppens/PromoterAtlas-data

## Code availability

The code associated with this study is available at https://github.com/LucasCoppens/PromoterAtlas This repository includes the final model weights of each of the models created in this work. In addition, it includes the code used for base model training as well as a logit plot visualisation script to generate logit plots like the ones shown in this study. The repository also includes a script for end-to-end whole-genome promoter annotation on genbank input file, as well as the resulting genbanks of whole-genome promoter annotation for a selection of medically and biotechnologically relevant Gammaproteobacteria studied in this work. Finally, it includes scripts that allow inference with this study’s expression prediction models and a script for distance calculation and prediction of co-regulated gene clusters.

## Acknowledgements

The authors thank Timon Schneider and Matthieu Meeus for insightful discussions that helped improve aspects of this work.

## Supplementary Figures

**Figure S1.**
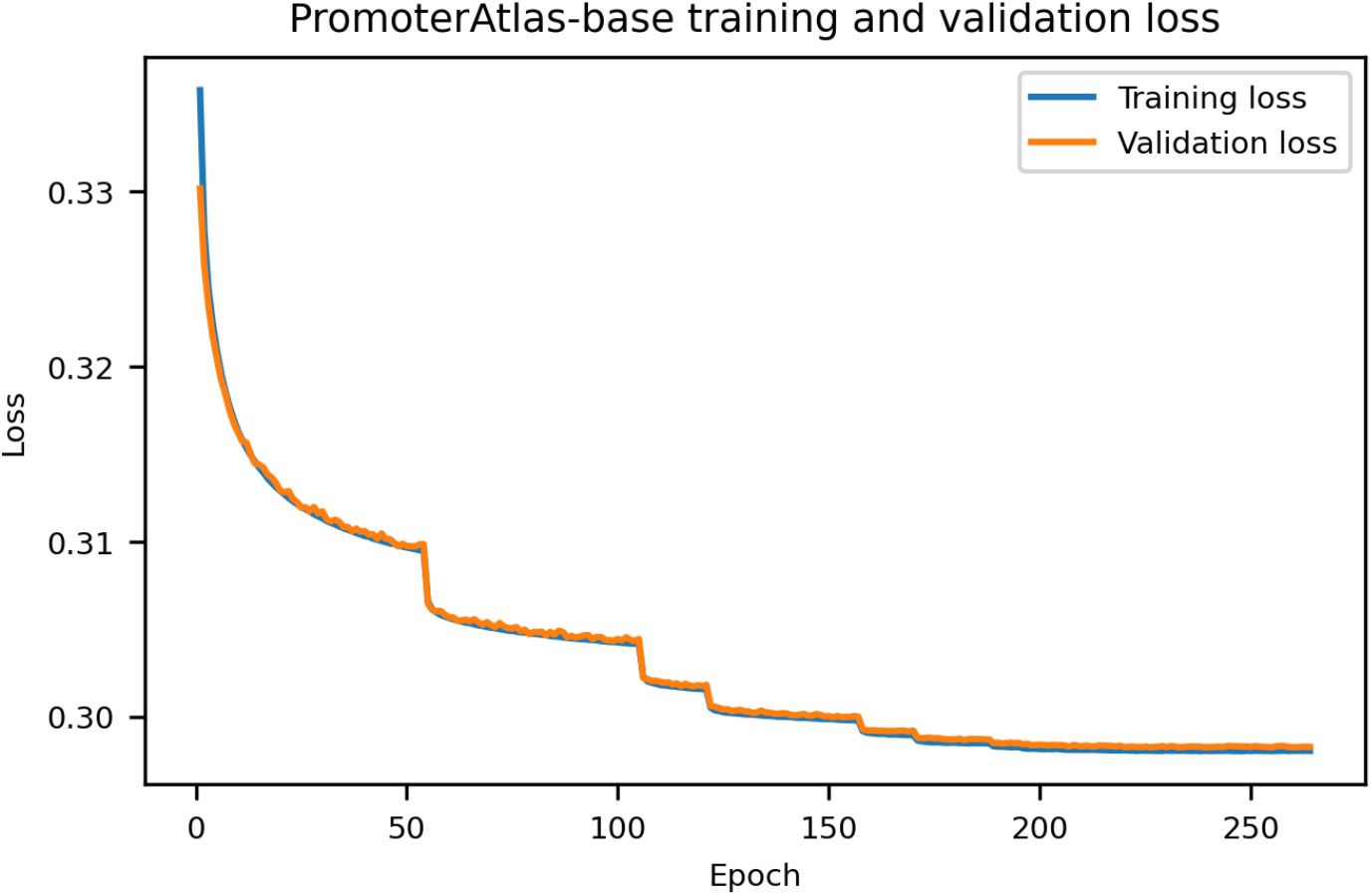
PromoterAtlas-base training loss plot. The curve shows good convergence of train and validation loss. The step-wise behaviour is explained by the learning rate being halved when a plateau is reached with the previous learning rate, allowing the model to learn finer detail each time.

**Figure S2.**
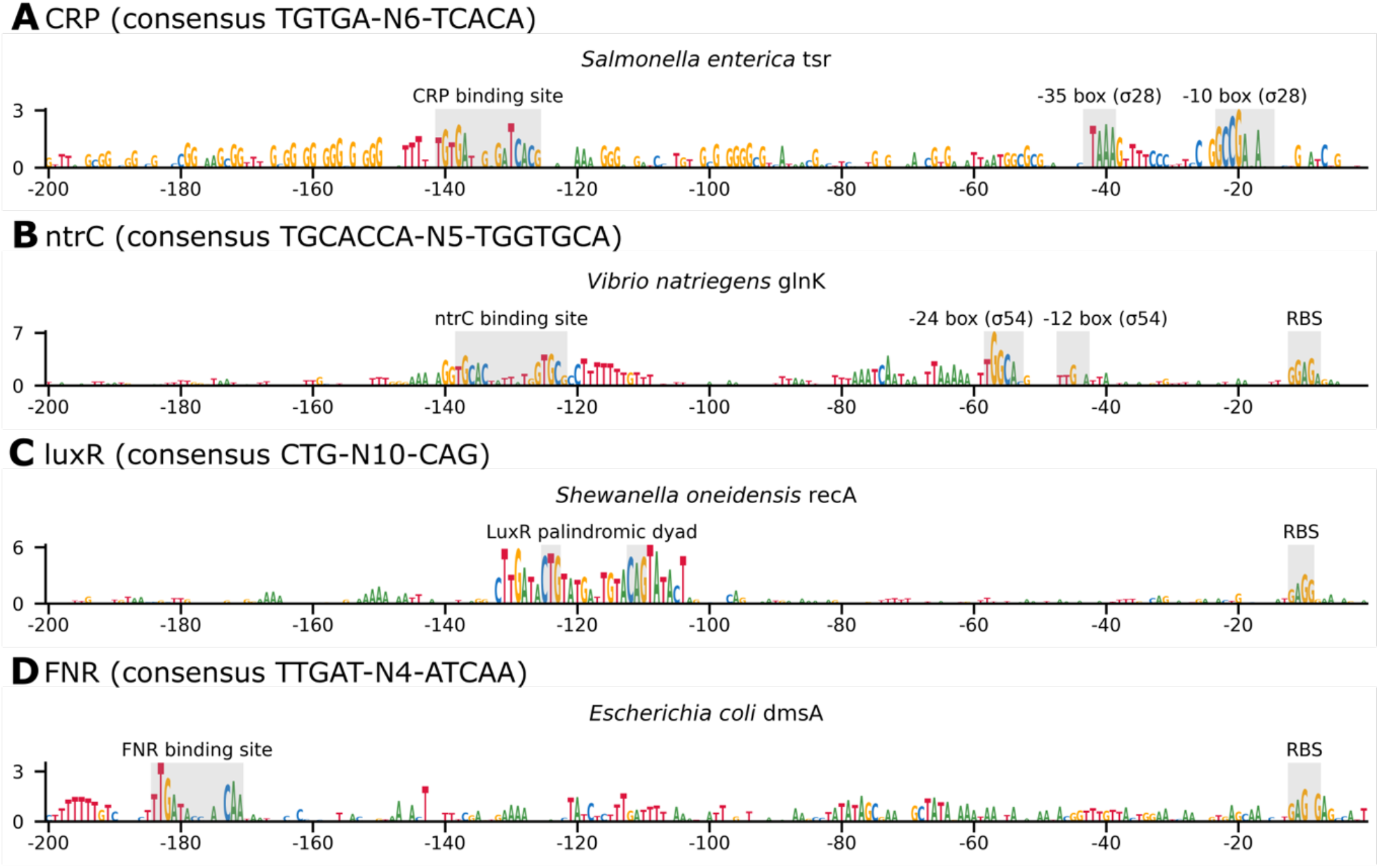
Examples of PromoterAtlas logit plots showing recognition of transcription factor binding site motifs for CRP, ntrC, luxR, and FNR.

**Figure S3.**
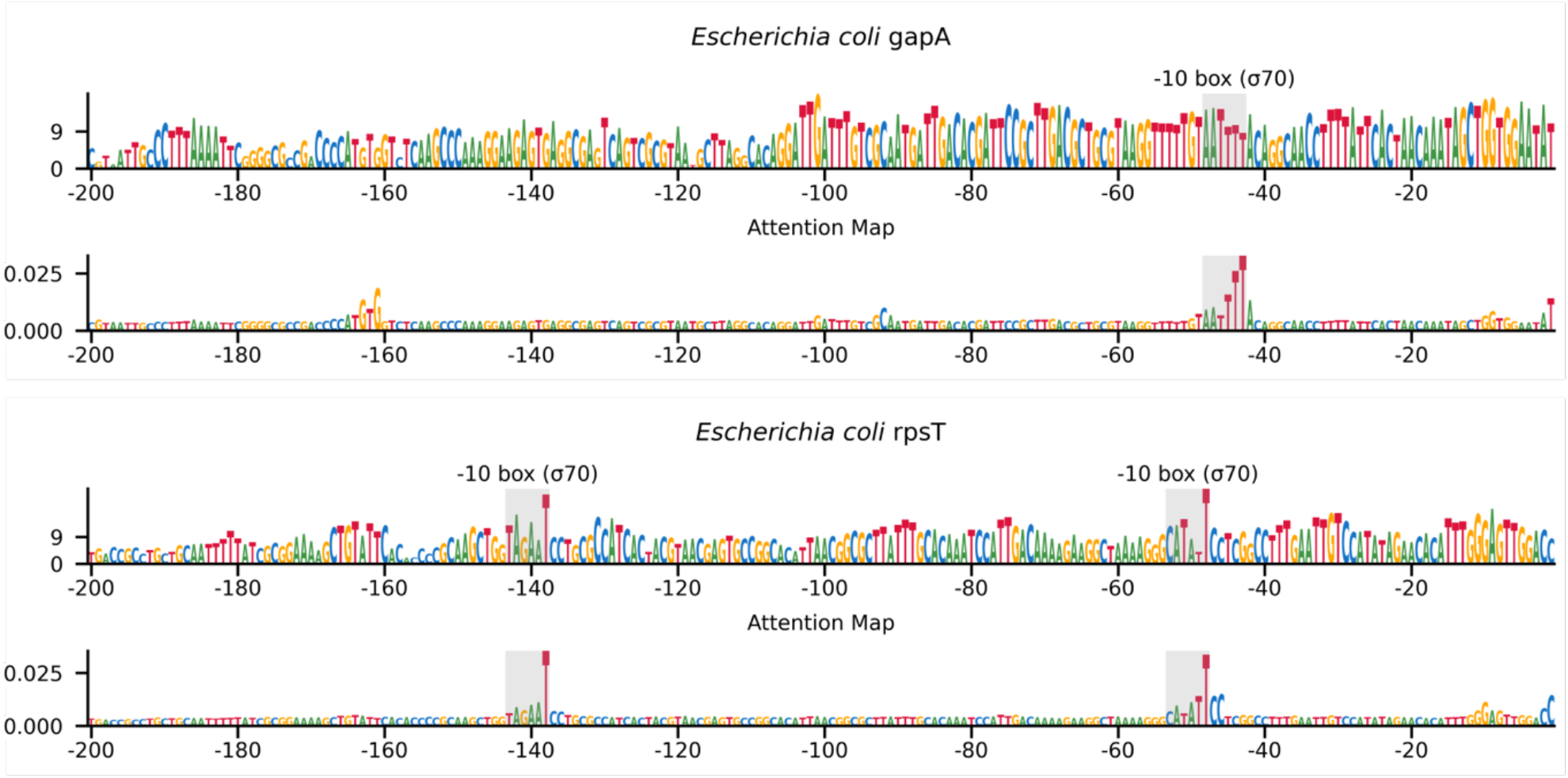
The regulatory regions upstream of some genes known to be highly expressed, long stretches of high logits in the plot prevent straightforward identification of regulatory elements. In those cases, we often found that plotting the PromoterAtlas’s average attention map would reveal the presence of σ70 motifs, exhibiting a peak in the logit values at the last position of the −10 motif box.

**Figure S4.**
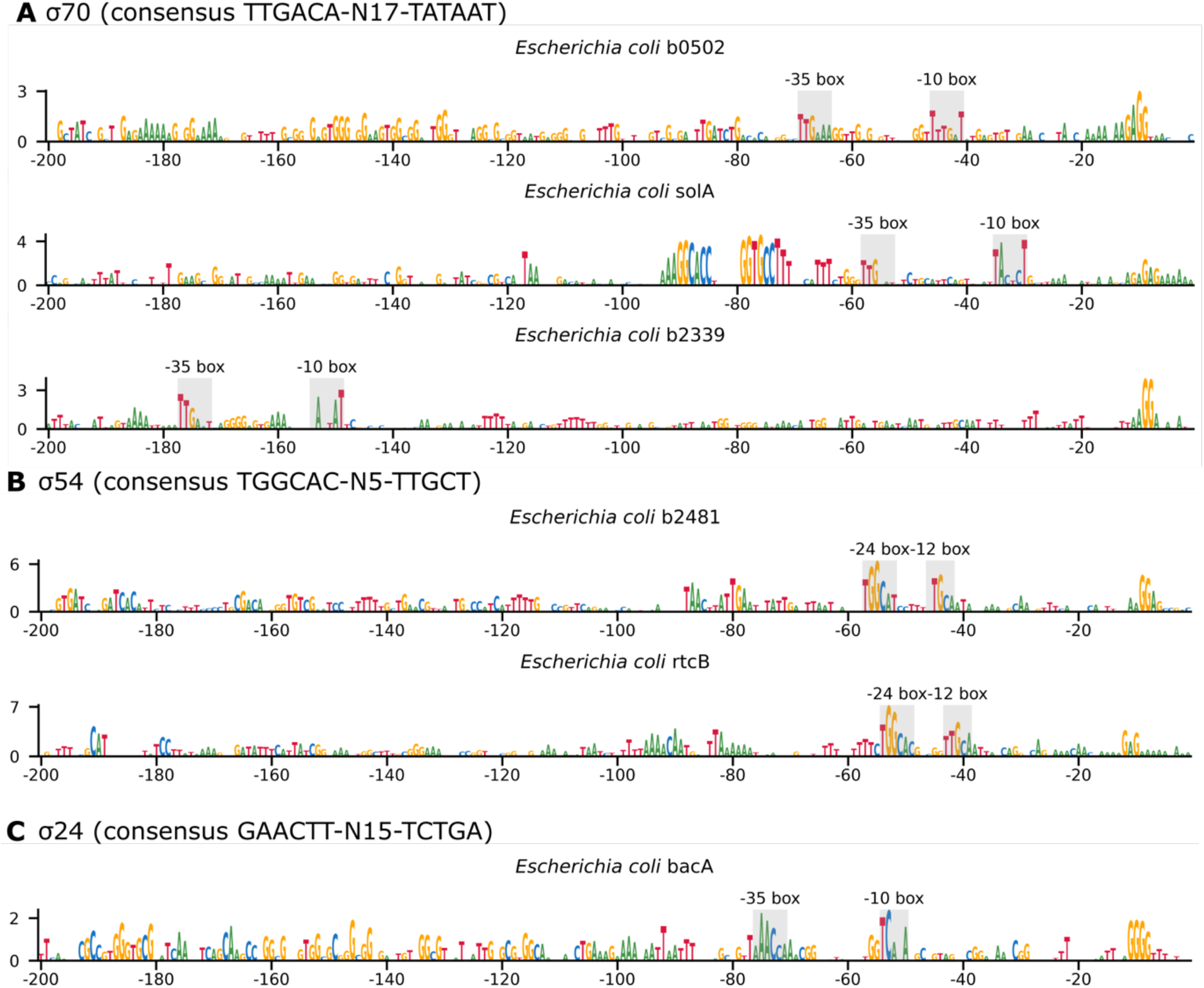
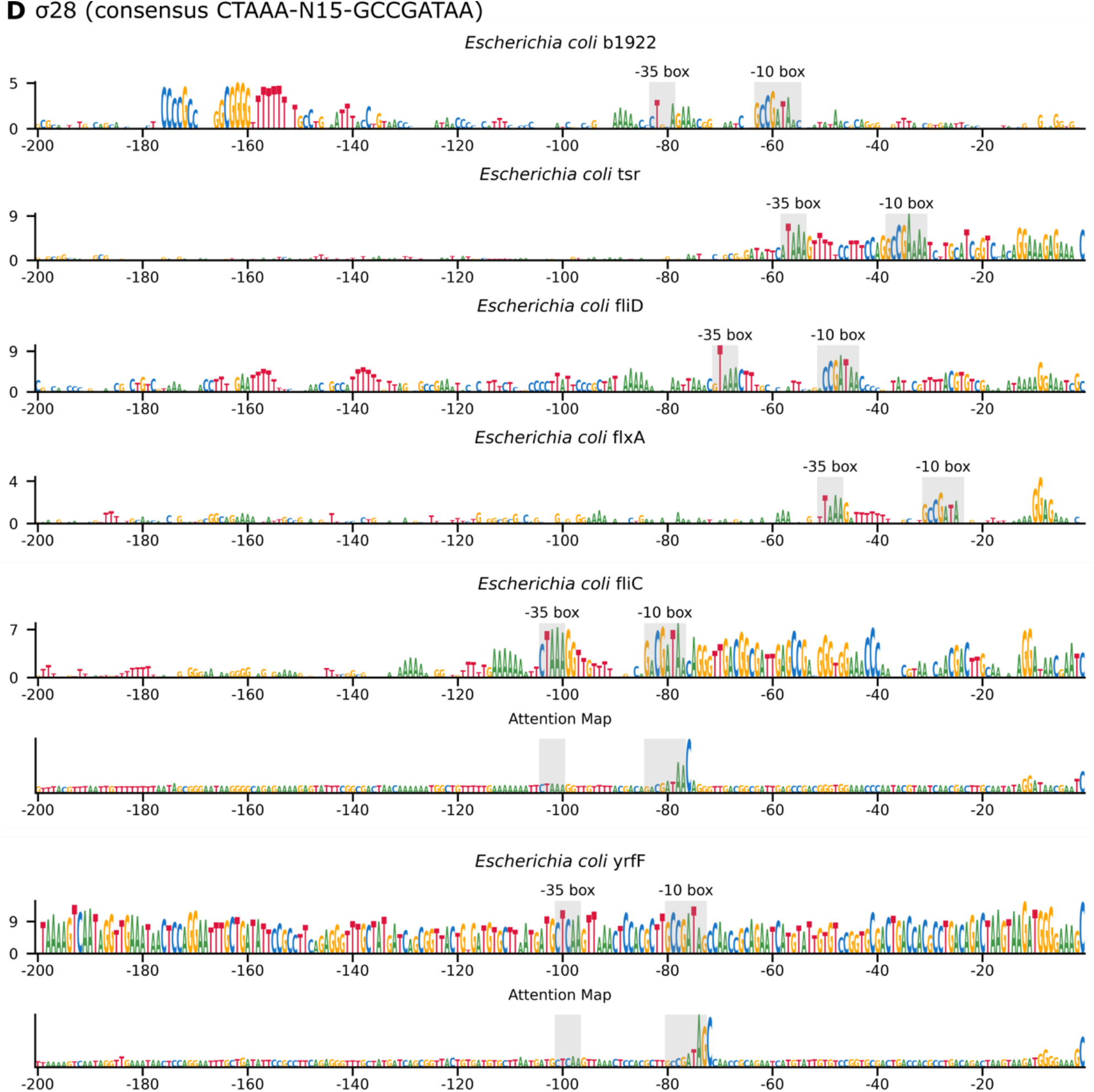
Logit plots of the *E. coli* promoters predicted by PromoterAtlas-segmentation but which did not match the Cho et al. (2014)^19^ ChIP-seq data show recognition of clear promoter motifs. (A) Logit plots of 3 σ70 promoters predicted by PromoterAtlas-segmentation but not validated by Cho et al. (2014)^19^. (B) Logit plots of the 2 σ54 promoters predicted by PromoterAtlas-segmentation but not validated by Cho et al. (2014)^19^. (C) Logit plots of the σ24 promoter predicted by PromoterAtlas-segmentation but not validated by Cho et al. (2014)^19^. (D) Logit plots of the 6 σ28 promoters predicted by PromoterAtlas-segmentation but not validated by Cho et al. (2014)^19^. For fliC and yrfF the attention maps are presented as well to show the positions of the −10 boxes as high logits prevent clear distinction in the logit plots.

**Figure S5.**
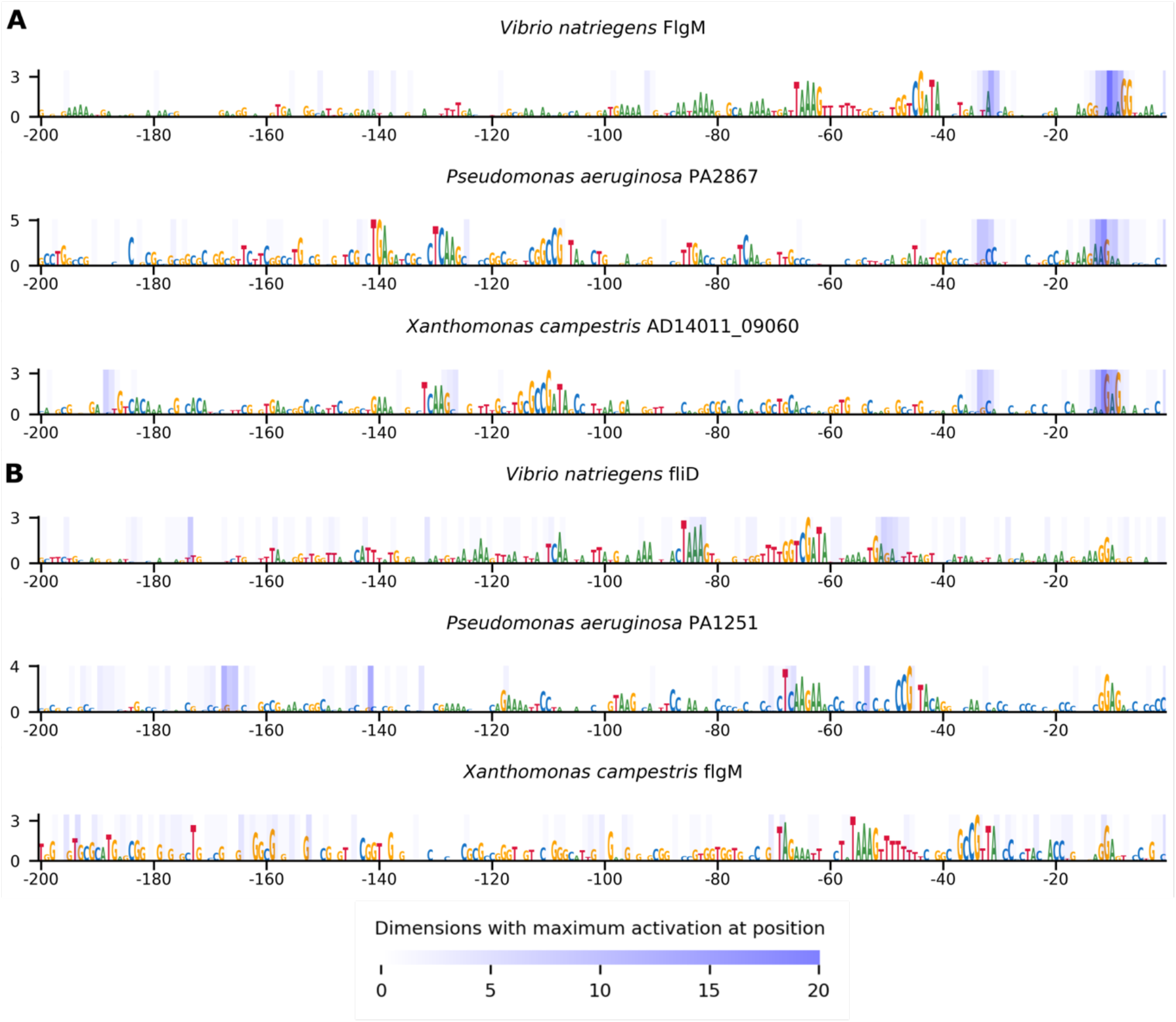
Visualisation of positions contributing most strongly to pooled sequence embeddings reveal two distinct encoding mechanisms for σ28 promoter sequences. (A) The embeddings of three σ28 promoter sequences from the σ28-exclusive cluster in the block 7 UMAP plot from Figure 4 are mostly determined by information encoded in the vicinities of the −10 and −33 positions, regardless of where the σ28 promoter motifs appear in the sequence. (B) Three examples of σ28 promoter sequences from the main heterogeneous cluster in the block 7 UMAP plot from Figure 4 which do not show the same characteristic encoding pattern as the σ28-specific cluster.

**Figure S6.**
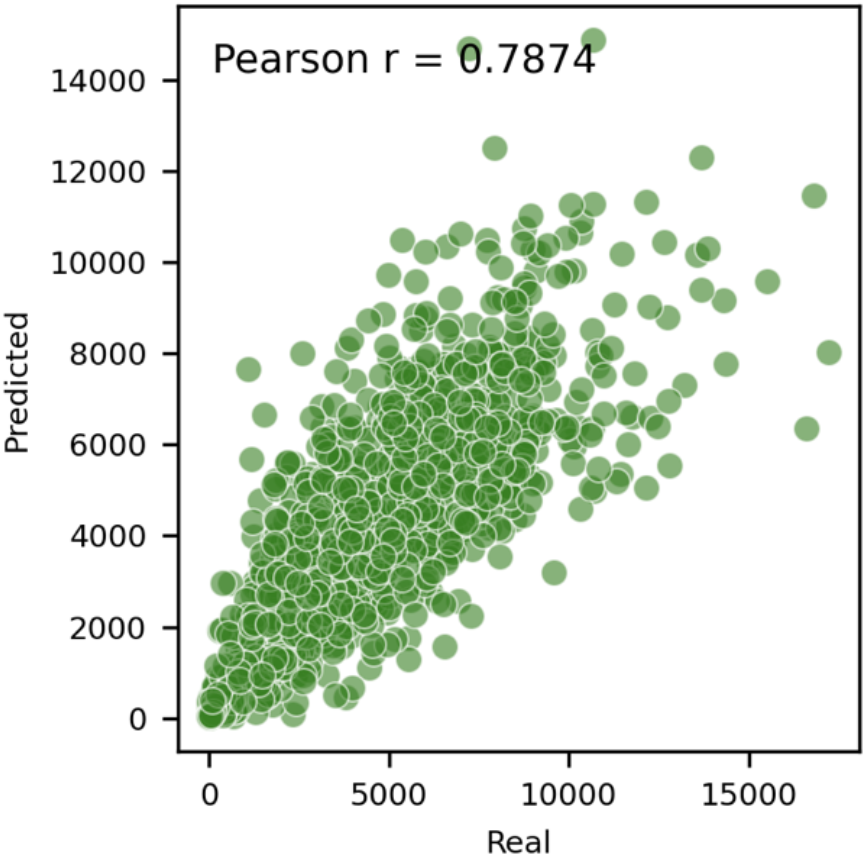
Scatter plot of real protein expression values from the Kosuri et al. (2013)^31^ dataset and the values predicted by the PromoterAtlas protein expression model on the test data.

## Notes

https://github.com/LucasCoppens/PromoterAtlas

https://huggingface.co/datasets/LCoppens/PromoterAtlas-data

### Competing Interest Statement

The authors have declared no competing interest.

### Summary of Updates

The caption of figure S4 contained an error which has now been rectified.

